# Quantitative isotope-labeled crosslinker proteomics reveals developmental variation in protein interactions and posttranslational modifications in *Diaphorina citri*, the citrus greening insect vector

**DOI:** 10.1101/2021.11.09.467860

**Authors:** John S. Ramsey, Xuefei Zhong, Surya Saha, Juan D. Chavez, Richard Johnson, Jaclyn E. Mahoney, Andrew Keller, Kathy Moulton, Lukas A. Mueller, David G. Hall, Michael J. MacCoss, James E. Bruce, Michelle Heck

## Abstract

Acquisition of the citrus greening bacterial pathogen, ‘*Candidatus* Liberibacter asiaticus’ (*C*Las) by Asian citrus psyllid (*Diaphorina citri*) nymphs is required efficient tree-to-tree transmission during the adult stage. Quantitative isotope-labeled protein interaction reporter (PIR) cross-linkers were used in parallel with protein quantification using spectral counting to quantify protein interactions within microbe-enriched cellular fractions of nymph and adult *D. citri*. Over 100 unique crosslinks were found between five insect histone proteins, and over 30% of these were more abundant in nymph compared to adult insects. Strikingly, some cross-links detected in *D. citri* proteins are conserved in cross-linking studies on human cells, suggesting these protein interaction topologies were present in the common ancestor (∼750MYA) or are subject to convergent evolution. Analysis of posttranslational modifications of crosslinked histones revealed the presence of acetylated and methylated lysine residues, which may impact psyllid chromatin structure and gene expression. Histone H3 peptides acetylated in the N terminal tail region were found to be more abundant in nymph compared to adult insects in two orthogonal proteomics methods. The insect life stage-specific histone posttranslational modifications and protein interactions represent physical evidence that metamorphosis is associated with changes in chromatin structure that regulate genome-wide transcriptional reprogramming.

## Introduction

Insect transmission of bacterial and viral pathogens is responsible for the spread of numerous agricultural pathogens that cause diseases of economic significance (1, 2). The Asian citrus psyllid (ACP, *Diaphorina citri*), is one of two psyllid species responsible for the worldwide spread of ‘*Candidatus* Liberibacter asiaticus’ (*C*Las), the gram negative, fastidious bacteria responsible for huanglongbing (HLB), also referred to as citrus greening disease. *C*las is a gram-negative α-proteobacteria which is not amenable to laboratory culture by standard methods, but which is capable of replicating in the disparate environments of citrus phloem sap and tissues of the insect vector (3, 4). Evidence on the mode of *C*Las transmission is consistent with a circulative, propagative transmission pathway in *D. citri*, whereby bacteria acquired into the gut of the insect vector from feeding on an infected tree must circulate and replicate within the insect body and enter the salivary glands in order to be transmitted to a healthy tree as a component of the insect’s saliva (5). While both nymph and adult insects are capable of acquiring *C*Las into their guts, the bacterium can only be transmitted by adult insects that have acquired the pathogen during their nymphal stage (6). The weaker nymphal immune response may enable the bacteria to readily cross the gut transmission barrier in nymphs, which is fortified and difficult to penetrate in adults (7).

Analysis of the role of genes, transcripts, proteins, and metabolites in *C*Las transmission by *D. citri* may reveal novel approaches to disease control (8, 9). High-throughput genome and transcriptome analysis has provided details of gene sequences and expression levels in a wide range of insect vectors (10-12). Proteomics analyses rely on databases of proteins predicted from genomic and transcriptomic data, and application of mass spectrometry workflows to the study of vector biology has enabled the identification of proteins and metabolites that are present at high abundance in specific transmission barrier tissues or in response to experimental treatments such as host plant infection (13-16). Mass spectrometry-based workflows designed to identify protein-protein interactions provide complementary data to quantitative proteomics studies, revealing details of protein intramolecular structure. The conformation of protein complexes which may enable the development of targeted strategies to block pathogen transmission by insect pests (17, 18).

Crosslinking mass spectrometry (XL-MS) using protein interaction reporter (PIR) technology has been used for *in vivo* identification of specific regions of interaction between protein domains and within protein complexes in cell and tissue samples (17, 19, 20). A prior study combining PIR and quantitative proteomics identified protein interactions between adult *D. citri* proteins and bacterial proteins. Quantitative proteome analysis of nymph and adult insects from *C*Las(-) and *C*Las(+) colonies revealed a much greater impact of infection status on adult as compared to nymphal-stage insects (18). The less robust immune response to *C*Las observed in nymphs compared to adults may be related to the fact that *C*Las acquisition by nymphs is critical to enabling subsequent transmission by adults. The number of differentially abundant proteins identified between nymph and adult insects was several-fold greater than between *C*Las(-) and *C*Las(+) colonies within either developmental stage (18). More than 2000 proteins (nearly half of all identified proteins) were identified by spectral counting as differentially abundant between nymph and adult insects, of which several hundred were undetected at one developmental stage or the other (18). This is consistent with prior studies documenting genome-wide transcriptional changes regulated by changes in chromatin accessibility and remodeling associated with insect metamorphosis (21). The PIR results from that study yielded only very few cross-linked peptides between *D. citri* and *C*Las proteins proteins (18). Peptides from *C*Las proteins predicted to catalyze the first two steps in the conversion of pantothenate to coenzyme A were found to be crosslinked to the *D. citri* proteins myosin and hemocyanin 1, the latter of which has extensively documented functions in arthropod metabolism and immunity (22-24).

In the current study, we attempted to increase the identification of microbe involved cross-linked peptides by using a percoll gradient centrifugation strategy to produce cellular fractions from *C*Las(+) nymph and adult *D. citri*. These fractions were treated with isotopic variant PIR crosslinkers developed for quantitative PIR analysis, identical except for eight deuterium/hydrogen substitutions, were used to separately crosslink adult and nymph samples that were subsequently combined into a single sample for combined mass spectrometry analysis (25). Quantitative analysis of differences in crosslink abundance between developmental stages was performed in biological and technical replicate samples. In parallel, quantitative proteomics workflows were used to analyze the same Percoll fractions from *C*Las(+) and *C*Las(-) adult and nymph insects, and statistical analysis of spectral counting data revealed proteins whose abundance varied with developmental stage and/or infection status. Together, these data types allow us to distinguish between the developmental regulation of protein interaction formation as a result of changes in protein abundance or because of a change in protein interaction topology and protein complex formation during insect development.

## Experimental Section

### Insect Rearing

A colony of *D. citri* was continuously reared on *C*Las-infected *Citrus medica* at the USDA Agricultural Research Service Horticultural Research Laboratory in Fort Pierce, Florida. for several generations. Monthly quantitative PCR (qPCR) tests were performed on to determine the percent infection and average *C*Las Cq values as described (5). *C*Las(+) adult and nymph psyllids were collected from colonies in which the percent infection was 80%, with an average Cq value of 27.5. QPCR was performed on triplicate Percoll gradient fractionations to assess the distribution of CLas titer in the fractionation scheme (**Table 1**). Analysis of the *D. citri* wingless gene was performed as described (26) to assess the distribution of *D. citri* cells in the fractions (**Table 1**).

**Table 1.** Average CLas and wingless gene quantification by qPCR in Percoll gradient fractions

| Fraction | Average CLas Cq<sup>2</sup> | Average Wg Cq<sup>2</sup> |
| --- | --- | --- |
| 1, 2 | 26.18 (0.17) | 23.36 (0.14) |
| 3,4 | 24.38 (0.07) | 24.43 (0.06) |
| 5,6 | 20.65 (0.04) | 21.33 (0.03) |
| 7,8 | 23.78 (0.08) | 24.46 (0.06) |
| 9,10 | 25.10 (0.06) | 26.80 (0.09) |
| 11,12 | 29.52 (0.07) | 29.69 (0.23) |
| Whole<sup>1</sup> | 19.29 (0.16) | 23.57 (0.07) |

<sup>1</sup> Average Cq values for pooled whole insects used prior to Percoll gradient fractionation.
<sup>2</sup> Standard deviation of the mean in parenthesis.

### Experiment Design and Statistical Rationale

A total of 8 insect samples [4 biological replicates each adult *C*Las(+) and nymph *C*Las(+) samples] were fractionated using Percoll density centrifugation and the *C*Las-enriched fractions from each replicate were pooled and analyzed using the Quantitative Protein Interaction Reporter (qPIR) workflow. Each biological replicate comprises a population of insects (500 adults per sample, 750 nymphs per sample) to capture the biological variation within insect populations. Four biological replicates for each sample type (nymph/adult) were used to account for variability in sample preparation (**Figure 1**). Triplicate injection replicates were performed to account for analytical variability in sampling PIR labeled peptides. For each peptide crosslink identified in pooled nymph/adult samples, the 95% confidence interval was calculated for the mean log2 ratio of nymph/adult crosslink abundance. A crosslink was identified as more abundant in nymph or adult samples when the {[absolute value of crosslink log2 ratio] – [1/2 (95% CI)]} ≥ 1.

**Figure 1.**
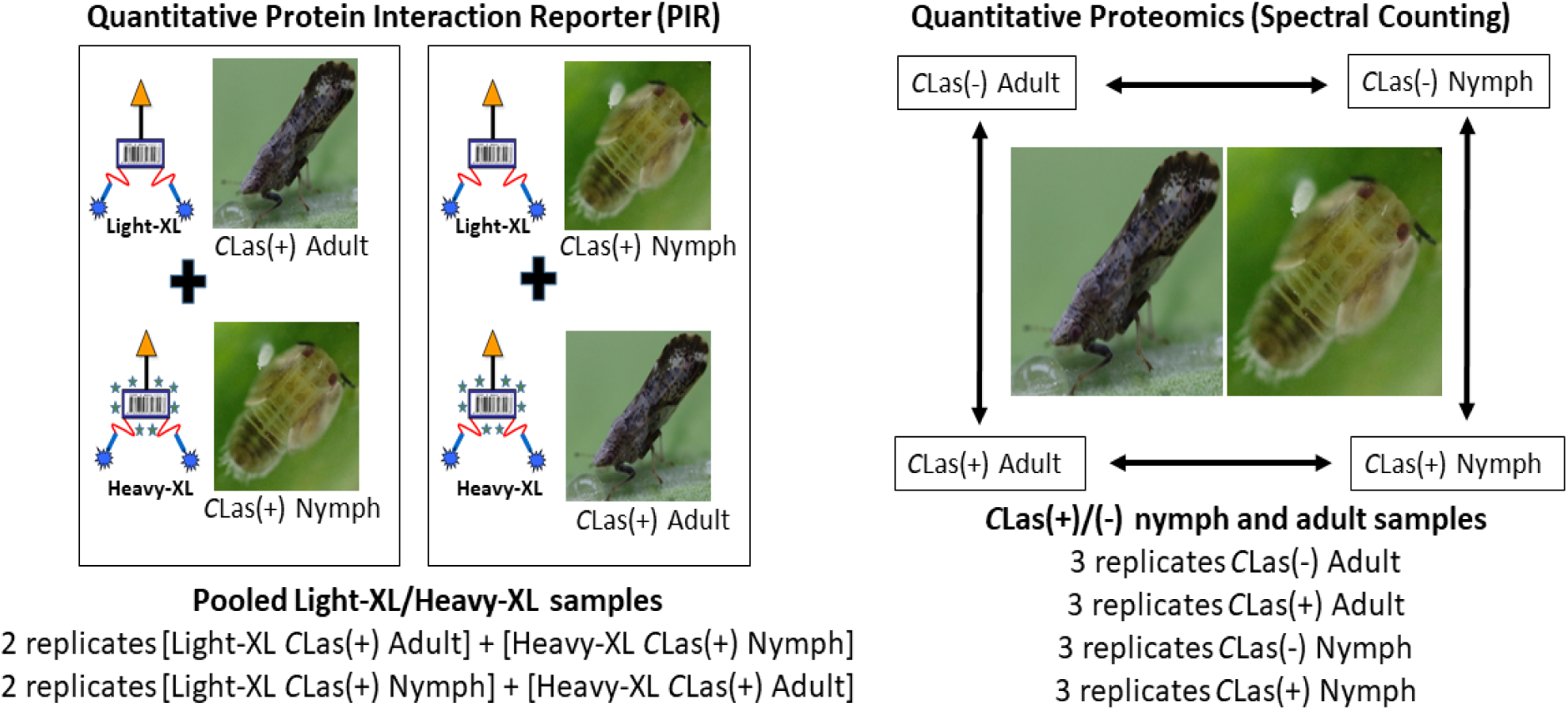
Experimental design of *Diaphorina citri* Quantitative Protein Interaction Reporter (PIR) and Quantitative Proteomics (Spectral Counting) studies. In the PIR study, percoll gradient cellular samples prepared from nymph and adult *D. citri* were treated with light or heavy crosslinkers (Light-XL, Heavy-XL) which generate unique mass profiles. Nymph and adult samples (four each) treated with Light-XL and Heavy-XL were pooled prior to peptide sample preparation and mass spectrometry analysis, and differences in crosslink abundance between nymph and adult samples were identified in replicate reciprocally labeled samples. In the Spectral Counting study, three replicates each of *C*Las(+)/*C*Las(-) nymph and adult percoll fraction samples were analyzed by mass spectrometry, and differentially abundant proteins were identified by pairwise statistical analysis between sample classes.

A total of 12 insect samples [3 biological replicates each adult *C*Las(+), adult *C*Las(-), nymph *C*Las(+), nymph *C*Las(-)] were taken through the Percoll fractionation and the same fractions as above were analyzed using data dependent acquisition and quantitative proteomics using spectral counting (Figure 1). The Fisher’s exact test using peptide spectral counting was used to identify differentially abundant proteins in pairwise analyses between the four sample classes. Benjamini-Hochberg p-value adjustment for multiple testing was applied to determine threshold for differential abundance.

### Quantitative Protein Interaction Reporter (PIR) sample preparation and analysis

Adult (∼500 insects per sample) and nymph (∼ 750 insects per sample) *D. citri* collected from *C*Las infected colonies were homogenized and fractionated by Percoll density centrifugation (14). Fractions 3, 4, 5 and 6 from each sample were pooled and suspended in a buffered solution consisting of 50 mM Na_2_HPO_4_ and 0.85 % (w/v) NaCl pH 7.4 for subsequent crosslinking reaction. Four adult samples and four nymph samples were crosslinked with 10 mM of light or heavy PIR cross-linker (Biotin Aspartate Proline-N-hydroxyphthalamide, BDP-NHP) at room temperature for 1 hour (25). Within each developmental group, two samples were crosslinked with light PIR crosslinker (d_0_-BDP-NHP) and two samples were crosslinked with heavy PIR crosslinker (d_8_-BDP-NHP). After the reaction, cells in the reaction mixture were lysed by incubation with equal volume of 8 M urea in 0.1 M Tris buffer overnight and ultrasonication. Protein concentration was determined by Bradford assay. Equimolar nymph light-XL and adult heavy-XL (and reciprocally labeled) protein samples were combined for a single injection into the mass spectrometer, enabling relative quantitation of crosslinks in different sample types within a single instrument run. The pooled protein lysates were reduced with 5 mM tris(2-carboxyethyl)phosphine (TCEP) for 30 minutes and alkylated with 10 mM iodoacetamide (IAA) for 45 minutes. Samples were diluted with 0.1 M ammonium bicarbonate buffer (pH=8) to urea concentration below 1 M, followed by overnight trypsin (Promega, WI) digestion at 37°C with 1:200 (enzyme: total protein) ratio. The digested mixtures were desalted with 1cc Sep-Pak C18 solid phase extraction (SPE) cartridges (Waters, MD), dried in vacuum centrifuge, and fractionated by strong cation exchange (SCX) HPLC (250 × 10 mm, Luna 5 μm diameter, 100 Å pore size particles, Phenomenex, Torrance, CA). The peptides mixture were eluted with a binary solvent gradient (Solvent A, 7 mM KH_2_PO_4_ (pH 2.6)/ 30% ACN, Solvent B, 7 mM KH_2_PO_4_ (pH 2.6)/350 mM KCl) at 1.5 mL/min mobile phase flow rate (0−2.5 min, 0% B; 2.5−7.5 min, 0−5% B; 7.5−47.5 min, 5−60% B; 47.5−67.5 min, 60−100% B; 67.5−77.5 min, 100% B). Five fractions of eluates (42.5-52.5min, 52.5-57.5 min, 57.5-62.5 min, 62.5-67.5min, 67.5-87.5 min) were collected, and further dried by vacuum centrifuge, resuspended in 5 mL 0.1 M NH_4_HCO_3_, and pH adjusted to ∼7.4. To enrich the crosslinked peptides, each collected fractions was incubated with ∼200 µL UltraLink monomeric avidin beads slurry (Thermo Pierce, Rockford, IL) for 30 min. The crosslinked peptide pairs captured by the beads were eluted with 0.5 mL of 70% ACN with 0.5% formic acid. The samples were dried and resuspended in 0.1% formic acid prior to nano LC−MS analysis.

Samples from each SCX fraction were analyzed in triplicate by a Waters NanoAcquity UPLC coupled to a Thermo Velos Fourier transform ion cyclotron resonance mass spectrometer implemented with Real-time Analysis for Cross-linked peptide Technology (ReACT) (27). The cross-linked peptide pairs were separated by a in-housed packed 40 cm × 75 μm C8 column (5 μm diameter, 100 Å pore size MichromMagic beads) with a 4 hour 10–30% B gradient (solvent A: 0.1% formic acid in water, solvent B: 0.1% formic acid in acetonitrile, flow rate 300 nL/min). High resolution full MS survey scan and CID MS/MS scans for ions baring more than 3 charges were acquired by ICR cells at 50 k resolving power at m/z 400, to examine cross-linked peptide pair candidates in real-time. The presence of reporter ion mass (m/z 752.4129) in the CID MS/MS spectra, along with fragment ion masses satisfying the PIR mass relationship (precursor mass = reporter ion mass + peptide α mass + peptide β mass) within 20 ppm of mass tolerance among the top 500 fragment ions in the MS/MS spectra suggests the precursor ion as a potential candidate of cross-linked peptide. Real time Analysis for Cross-linked peptide Technology (ReACT) triggers subsequent low resolution CID MS/MS/MS scans of two linear peptides released from the cross-linked peptide pair candidate ion ‘on-the-fly’ (27). For each released peptide ion from one cross-linked peptide pair precursor, two MS/MS/MS scans were acquired by the linear ion trap with normalized collision energy of 35.

### PIR Data analysis

A database consisting of 4439 *Diaphorina citri* protein sequences (ftp://ftp.citrusgreening.org/manuscripts/Ramsey2020/PIR_database.fasta) identified from DDA-MS of *C*Las infected adult and nymph insects, 1012 *Candidatus Liberibacter asiaticus* sequences, 1032 *Wolbachia* sequences, 352 *Profftella* sequence, and 196 *Carsonella* sequences was used for identification of linear peptide sequences released from cross-linked peptide pairs. MS^3^ spectra from ReACT analysis were searched against the database containing both 7031 forward and 7031 reverse protein sequences by Comet (version 2017.01 rev. 4). The linear peptide sequences were then assembled to cross-linked peptide pairs according to the mass relationships recorded during the ReACT data acquisition. Comet search was conducted with the following parameters: precursor peptide mass tolerance 20 ppm, allowing for −1, 0, + 1, + 2, or +3 ^13^C offsets; fragment ion mass tolerance 1.005 Da with 0.4 Da offset; static modification, carbamidomethylation of Cys (57.0215 Da); variable modifications including Met oxidation (15.9949 Da), BDP stump modification (197.0324 Da for d_0_-BDP-NHP, 201.0597 Da for d_8_-BDP-NHP) of Lys and protein N-termini, where ‘stump’ refers to the residual group from the crosslinker attached to lysine side chain or peptide N-terminus after CID cleavage, Lys and Arg methylation (14.0156 Da), Lys dimethylation (28.0313 Da), Lys trimethsylation (42.0470 Da), Lys acetylation (42.0106 Da); up to three missed tryptic cleavages allowed. All the target and decoy MS^3^ pepetide-spectrum-matches containing an internal Lys modified by BDP stump were mapped back to the cross-linked relationships with a 20 ppm tolerance. XLinkProphet (28) was applied to model crosslink specific discriminating information, and the results were filtered at 1% FDR of unique crosslinks corresponding to minimum composite probability 0.63.

MasschroQ (29) was used for MS^1^ peak area integration of all crosslinked peptide pairs identified. LC-MS/MS runs acquired from the same SCX fractions were grouped for retention time alignment using the Obiwarp method. Accurate precursor masses of the most abundant three isotopes, charge state, and MS^2^ scan number of each peptide pair-spectra-match (PSM) of crosslinked peptide pair were used as input for MassChroQ. Peak areas under curves from extracted ion chromatograms were integrated for all input PSMs with 20 ppm mass tolerance of precursor ion using the Zivy peak detection algorithm (29). Log_2_(adult/nymph) ratio for each isotope of each PSM was alculated from peak area output by MassChroQ (29), and normalized by the median ratio within each biological replicates. All ratios belonging to one non-redundant crosslinked pair were grouped, the average ratio after outlier rejection was assigned as the ratio of the specific crosslinked peptide pair. The 95% confidence interval was computed for each non-redundant crosslink.

### Quantitative proteomics sample preparation and analysis

Three biological replicate samples of adult and nymph *D. citri* (∼ 200 insects per sample) collected from *C*Las-infected colonies were homogenized and fractionated by Percoll density centrifugation (14). The same four fractions collected for PIR analysis were used for protein extraction and peptide sample preparation, with fractions 3/4 and fractions 5/6 prepared processed separately for each biological sample. Protein precipitation solvent (10% trichloroacetic acid in acetone with 2% beta-mercaptoethanol) was made fresh and kept on ice until use. For Percoll gradient fraction protein extraction, one mL precipitation solvent was added to purified cells and the sample was sonicated to rupture cells. Several rounds of sonication (30 seconds, 15% amplitude, Branson digital probe sonicator, Danbury, CT) were needed to fully disrupt pellet, with samples chilled on ice between rounds of sonication to avoid heating. Protein samples were precipitated overnight in the -20°C freezer. Precipitated protein pellets were washed three times with one mL ice-cold acetone, and after decanting the supernatant from the final wash the pellet was dried to completion. Pellets were resuspended in 200 μL protein reconstitution solvent [8 M urea, 50 mM triethylammonium bicarbonate (TEAB) in water]. Pellets were reconstituted overnight with shaking. Samples were centrifuged at full speed to pellet insoluble material, and the supernatant was collected for protein analysis. Quick Start Bradford Protein Assay (Bio-Rad, Hercules, CA) was used to determine concentration of protein samples. Based on Bradford assay data, 3.5 μg of protein from Percoll gradient fraction samples was used for peptide sample preparation. For reduction, a 100 μL solution of each protein sample was prepared in 10 mM dithiotreitol, 50 mM TEAB, and samples were vortexed, centrifuged briefly, and incubated for one hour at 30°C. After cooling samples to room temperature, methyl methanethiosulfonate (MMTS) in 50 mM TEAB was added to a final concentration of 30 mM MMTS, samples were vortexed, centrifuged, and incubated for one hour at room temperature in the dark. Samples were diluted with TEAB if necessary to reduce concentration of urea below 1 M. Sequencing grade, modified trypsin (Promega, Madison, WI) was added to protein samples (target ratio of trypsin: protein is between 1:20-1:100 by weight). Samples were vortexed, centrifuged, and incubated at 30°C overnight.

Oasis MCX solid phase extraction cartridges (Waters, Milford, MA), 30 μM particle size, were used to remove contaminants from the digested peptides. Trypsin-digested peptide samples were dried in a vacuum concentrator and reconstituted in 0.1% formic acid in water. Concentrated formic acid (5 μL) was added to peptide samples, which were spotted on litmus paper to confirm pH>3. Cartridges were conditioned with 1) 1 mL 100% methanol, 2) 1 mL 3% ammonium hydroxide in water, 3) 2 mL 100% methanol, 4) 3 mL 0.1% formic acid in water. Samples were added to the column, salts were washed with 1 mL 0.1% formic acid in water, neutrals were washed with 1 mL 0.1% formic acid in methanol, and peptides were eluted in 1 mL 3% ammonium hydroxide in methanol and dried down in a speed-vac.

Mass spectrometry was performed on a Fusion (Thermo Fisher Scientific) mass spectrometer. Judging from the mass spectral response and total protein estimates, about one microgram of each sample digest was loaded from the autosampler onto a 150-μm Kasil fritted trap packed with Reprosil-Pur C18-AQ (3-μm bead diameter, Dr. Maisch) to a bed length of 2 cm at a flow rate of 2 ul/min. After loading and desalting using a total volume of 10 ul of 0.1% formic acid plus 2% acetonitrile, the trap was brought on-line with a pulled fused-silica capillary tip (75-μm i.d.) packed with the same Reprosil C18-AQ that was mounted in an in-house constructed microspray source and placed in line with a Waters Nanoacquity binary UPLC pump plus autosampler. Peptides were eluted off the column using a gradient of 2-25% acetonitrile in 0.1% formic acid over 100 minutes, followed by 25-60% acetonitrile over 40 minutes at a flow rate of 250 nl/min.

The mass spectrometer was operated using electrospray ionization (2 kV) with the heated transfer tube at 275 C using data dependent acquisition (DDA) in the so-called “Top Speed” mode, whereby one orbitrap mass spectrum (m/z 400-1600 with quadrupole isolation) was acquired with multiple linear ion trap tandem mass spectra every three seconds or less. The resolution for MS in the orbitrap was 120,000 at m/z 200, and for MS/MS the linear ion trap provided unit resolution. The automatic gain control targets for MS in the orbitrap was 2e5, whereas for MS/MS it was 1e4, and the maximum fill times were 20 and 35 msec, respectively. The MS/MS spectra were acquired using quadrupole isolation with an isolation width of 1.6 m/z and HCD collision energy (NCE) of 30%. The precursor ion threshold intensity was set to 1e4 in order to trigger an MS/MS acquisition. Furthermore, MS/MS acquisitions were allowed for precursor charge states of 2-7. Dynamic exclusion (including all isotope peaks) was set for 5 seconds using monoisotopic precursor selection with a mass error of 15 ppm. The fragment ions were analyzed in the linear trap using the “rapid” scan rate.

Mass spectrometry data files were searched against a combined database of predicted *D. citri, C*Las (GCF_000023765.2), and endosymbiont [‘*Candidatus* Carsonella ruddii’ DC (GCF_000441575.1), ‘*Candidatus* Profftella armatura’ (GCF_000441555.1), and *Wolbachia pipientis* (strain Wolbachia wDi, GCF_000331595.1], proteins using Mascot Daemon 2.5.1 (Matrix Science, Boston, MA). *D. citri* proteins from NCBI GNOMON annotation 100 of the diaci version 1.1 genome and MCOT transcriptome (30) were clustered with cd-hit (31) to remove redundancy (parameters: -c 1.00 -d 0 -n 5). The resultant *D. citri* protein set was merged with Carsonella, Profftella, *Wolbachia*, and *C*Las to create the combined database (ftp://ftp.citrusgreening.org/manuscripts/Ramsey2020/Gnomon_MCOTahrd_clustered_endosymbionts.faa). MS/MS search parameters included fixed modifications (cysteine: methylthio), variable modifications (lysine: methylation, dimethylation, acetylation; arginine: acetylation; asparagine, glutamine: deamidation; methionine: oxidation), and maximum one missed cleavage. Thermo *.raw files were converted into Mascot Generic Format using MSConvert in ProteoWizard (32). Files with the *.dat extension were exported from Mascot and loaded into Scaffold Q+ 4.8.9 (Proteome Software, Portland, OR) and used to calculate normalized spectral count for each protein from each sample. Scaffold protein and peptide thresholds were set at 95%, with a minimum peptide number of two per protein – this resulted in a protein false discovery rate (FDR) of 1.8% and a peptide FDR of less than 0.1% (33, 34). Statistical analysis of weighted spectral count data for identification of proteins differentially abundant between sample categories was performed in Scaffold using Fisher’s exact test (significance level p<0.05 using the Benjamini-Hochberg multiple test correction). The mass spectrometry proteomics data have been deposited to the ProteomeXchange Consortium via the PRIDE partner repository with the dataset identifier PXD020412 (35, 36).

### Protein structure modeling

ICM (Molsoft LLC, San Diego, CA) and Rasmol (37) software were used to visualize .pdb files representing predicted protein three dimensional structure.

## Results and Discussion

### Protein Interaction Reporter crosslink quantification

A *D. citri* protein interaction network was established using isotope-labeled Protein Interaction Reporter (PIR) mass spectrometry of microbe-enriched cellular fractions isolated from adult and nymph *D. citri* collected from *C*Las-infected citrus trees. QPCR analysis on three Percoll gradient fractionations showed *C*Las found in every fraction and slightly enriched in fractions 3 and 4 (**Table 1**). This crosslinking network is available at XLinkDB 3.0 http://xlinkdb.gs.washington.edu/xlinkdb/index.php (network name: ACP_Percoll_Fraction_XL_Bruce), which provides visualization of the quantitative site interaction network and details of identified crosslinks (38, 39). A total of 1571 non-redundant crosslinked peptide pairs were identified (**Table S1**), including 326 crosslinks between peptides derived from 93 unique protein pairs (**Table S2**). Out of the total 1571 crosslinks identified, 665 were quantified in two or more biological replicates from adult and nymph samples – for these, log2 ratios [with 95% confidence interval (CI)] of crosslink abundance between nymph and adult samples were calculated. Crosslinks were determined to be more abundant in one sample type (nymph or adult) when {[absolute value of crosslink log2 ratio] – [1/2 (95% CI)]} ≥ 1. Using these criteria, 78 crosslinks involving peptides from a total of 24 proteins were determined to be more abundant in adult *D. citri* compared to nymph *D. citri*, and 162 crosslinks involving peptides from a total of 82 proteins were determined to be more abundant in nymph *D. citri* compared to adult *D. citri* (**Table S3**).

### Functional characterization of differentially abundant crosslinks

These differentially abundant crosslinks were assigned to functional categories according to the annotation of the proteins associated with the crosslinked peptides (**Figure 2**). Crosslinks within and between a small number of myosin proteins comprise a large fraction of the total identified crosslinks, and the protein functional category most represented in the crosslinks found to be more abundant in both nymph and adult samples is cytoskeleton/muscle (**Figure 2**). This indicates that unique conformations or complexes of cytoskeleton/muscle proteins are present in nymph and adult insects, which is likely related to the dramatic differences in morphology and behavior between developmental stages, including the presence of wings and the concomitant significance of flight muscle in adults. Histone proteins are the second most represented category of crosslinks found to be more abundant in nymph *D. citri*, while no crosslinks containing histone proteins were more abundant in adult *D. citri* (**Figure 2**). A total of 109 unique crosslinks containing peptides mapping to histone proteins were identified, and 35 of these histone crosslinks were found to be more abundant in the nymph samples (**Table S1, S3**). Out of these 35 differentially abundant crosslinks, 12 are intermolecular crosslinks representing interactions between five pairs of histones: Histone 3/4; Histone 2b/4; Histone 3/2b; Histone 3/2a; Histone 2a/2b (**Table 2**).

**Table 2.** Histone cross-linked peptides detected as higher in relative abundance in *C*Las (+) nymphs vs. adult insects

| Peptide A<sup>1</sup> | Peptide B<sup>1</sup> | Protein A accession and description | Protein B accession and description | Crosslink log2ratio adult/nymph |
| --- | --- | --- | --- | --- |
| TVTAMDVVYALK<sup>118</sup>R | YQK<sup>57</sup>STELLIR | MCOT08963.0.MT_Histone H4 | MCOT05881.0.CC_Histone H3 | -4.83 |
| DAVTYTEHAK<sup>104</sup>R | EIAQDFK<sup>80</sup>TDLR | MCOT08963.0.MT_Histone H4 | MCOT05881.0.CC_Histone H3 | -2.57 |
| GVLK<sup>86</sup>VFLENVIR | LLPGELAK<sup>109</sup>H AVSEGTK | MCOT08963.0.MT_Histone H4 | A0A1S3DIC0_Histone H2B | -2.39 |
| DAVTYTEHAK<sup>104</sup>R | LLPGELAK<sup>109</sup>H AVSEGTK | MCOT08963.0.MT_Histone H4 | A0A1S3DIC0_Histone H2B | -2.07 |
| DAVTYTEHAK<sup>104</sup>R | K<sup>35</sup>ESYAIYIK | MCOT08963.0.MT_Histone H4 | A0A1S3DIC0_Histone H2B | -3.49 |
| YQK<sup>57</sup>STELLIR | K<sup>139</sup>SETAK | MCOT05881.0.CC_Histone H3 | MCOT13323.0.CO_Histone H2A | -2.11 |
| VTIMPK<sup>123</sup>DIQLAR | LLSGVTIAQGG VLPNIQAVLLP K<sup>144</sup>KTEK | MCOT05881.0.CC_Histone H3 | MCOT08626.0.MT_Histone H2A | -2.77 |
| YQK<sup>57</sup>STELLIR | LLSGVTIAQGG VLPNIQAVLLP K<sup>144</sup>KTEK | MCOT05881.0.CC_Histone H3 | MCOT08626.0.MT_Histone H2A | -3.93 |
| K<sup>65</sup>LPFQR | K<sup>35</sup>ESYAIYIK | MCOT05881.0.CC_Histone H3 | A0A1S3DIC0_Histone H2B | -2.85 |
| EIAQDFK<sup>80</sup>TDLR | LLPGELAK<sup>109</sup>H AVSEGTK | MCOT05881.0.CC_Histone H3 | A0A1S3DIC0_Histone H2B | -2.74 |
| K<sup>62</sup>GNYAER | K<sup>35</sup>ESYAIYIK | MCOT08626.0.MT_Histone H2A | A0A1S3DIC0_Histone H2B | -3.12 |
| K<sup>62</sup>GNYAER | ESYAIYIK<sup>44</sup>VLK | MCOT08626.0.MT_Histone H2A | A0A1S3DIC0_Histone H2B | -3.7 |

<sup>1</sup> Crosslinked residues are indicated in bold with crosslinked residue number in superscript.

**Figure 2.**
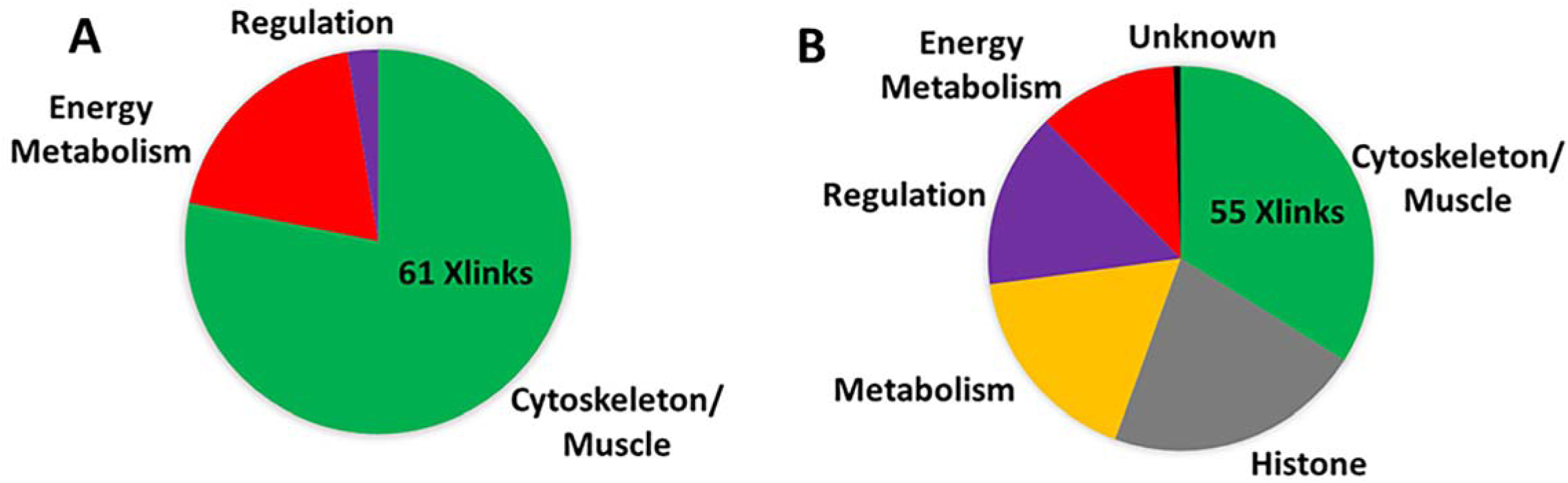
Functional categories assigned to proteins involved in crosslinks found to be more abundant in **a)** adult *D. citri* samples, and **b)** nymph *D. citri* samples. For each sample type, the number of crosslinks associated with 17 the most represented category is given. Detailed information on differentially abundant crosslinks is given in **Tables S5**. The category entitled Regulation included proteins involved in transcription, translation, and posttranslational modification.

In the remaining 23 differentially abundant histone crosslinks, both peptides map to the same protein (**Table 3**). Eight of these 23 crosslinks included peptides from histone H3 with posttranslational modifications (PTMs, one dimethylation and seven acetylations). Two of these histone H3 crosslinks include a peptide acetylated at K^10^ and a third includes a peptide dimethylated at the same position. In contrast to the convention in our predicted protein database, the histone numbering convention used in the literature counts the amino acid after the initiator methionine as the first residue – on account of this, K^10^ in our histone H3 protein is universally described in the literature as H3K9. Both acetylation and methylation of H3K9 are known to increase the hydrophobicity of the histone H3 tail, with the consequence of rendering this domain more available for binding of proteins functioning in chromatin remodeling, including chromodomain helicase DNA binding protein 3 (CHD3) (40). One of the histone crosslinks found to be more abundant in nymphs is the histone H3 homodimer crosslinked at H3K4 (T**K**^5^QTAR, **Table 3**). The intranucleosomal distances between the crosslinked residues of histone H3 protein pairs were estimated from analysis of the .pdb structure (PDB 1KX5) of the published X-ray crystal structure of the nucleosome(41). In cases where the predicted intranucleosomal distance between crosslinked homodimer residues was greater than the length of the crosslinker (30 angstroms), these crosslinks likely represent internucleosomal interactions. The lysine residues involved in the histone H3 crosslinks from **Table 3** are depicted on structural representations of the two copies of histone H3 within the nucleosome octamer (**Figure 3)**.

**Table 3.** Crosslinks where both peptides are derived from the same histone protein which are more abundant in nymph compared to adult *D. citri*.

| Peptide A | Peptide B<sup>1,2</sup> | PTM<br>(Peptide A<br>or B) | Protein accession and<br>description | Crosslink<br>log2ratio<br>adult/nym<br>ph |
| --- | --- | --- | --- | --- |
| TK<sup>5</sup>QTAR | K<sup>10</sup><sub>Me2</sub>STGGK<sup>15</sup>APR | Dimethylati<br>on (B) | MCOT05881.0.CC_Histo<br>ne H3 | -2.12 |
| K<sup>10</sup><sub>Ac</sub>STGGK<sup>15</sup>A<br>PR | K<sup>19</sup>QLATK<sup>24</sup><sub>Ac</sub>AAR | Acetylation<br>(A and B) | MCOT05881.0.CC_Histo<br>ne H3 | -3.5 |
| K<sup>10</sup>STGGK<sub>Ac</sub>AP<br>R | K<sup>19</sup>QLATK<sup>24</sup><sub>Ac</sub>AAR | Acetylation<br>(A and B) | MCOT05881.0.CC_Histo<br>ne H3 | -3.14 |
| K<sup>19</sup>QLATK<sup>24</sup><sub>Ac</sub>A | YQK<sup>57</sup>STELLIR | Acetylation | MCOT05881.0.CC_Histo | -3.14 |
| TK<sup>5</sup>QTAR | K<sup>10</sup>STGGK<sup>15</sup><sub>Ac</sub>APR | Acetylation | MCOT05881.0.CC_Histo | -3.03 |
| TK<sup>5</sup>QTAR | K<sup>19</sup>QLATK<sup>24</sup><sub>Ac</sub>AAR | Acetylation | MCOT05881.0.CC_Histo | -3.07 |
| TK<sup>5</sup>QTAR | K<sup>10</sup><sub>Ac</sub>STGGK<sup>15</sup>APR | Acetylation | MCOT05881.0.CC_Histo | -2.73 |
| TK<sup>5</sup>QTAR | K<sup>19</sup><sub>Ac</sub>QLATK<sup>24</sup>AAR | Acetylation | MCOT05881.0.CC_Histo | -2.79 |
| K<sup>65</sup>LFPQR | YQK<sup>57</sup>STELLIR | N/A | MCOT05881.0.CC_Histo | -3.75 |
| K<sup>10</sup>STGGK | TK<sup>5</sup>QTAR | N/A | MCOT05881.0.CC_Histo | -3.15 |
| TK<sup>5</sup>QTAR | YQK<sup>57</sup>STELLIR | N/A | MCOT05881.0.CC_Histo | -2.9 |
| TK<sup>5</sup>QTAR | STGGK<sup>15</sup>APR | N/A | MCOT05881.0.CC_Histo | -2.95 |
| TK<sup>5</sup>QTAR | TK<sup>5</sup>QTAR | N/A | MCOT05881.0.CC_Histo | -2.45 |
| K<sup>88</sup>YLVSAAVAA<br>GK | VDAQK<sup>81</sup>LAPFIR | N/A | MCOT07573.0.MM_Hist<br>one H1.1 | -2.73 |
| TEK<sup>148</sup>KA | LLSGVTIAQGGVLPNIQAVLLP<br>K<sup>144</sup>K | N/A | MCOT08626.0.MT_Hist<br>one H2A | -3.48 |
| LAHYNK<sup>86</sup>R | K<sup>35</sup>ESYAIYIK | N/A | A0A1S3DIC0_Histone | -3.4 |
| RK<sup>35</sup>ESYAIYIY | VLK<sup>47</sup>QVHPDTGVSSK | N/A | A0A1S3DIC0_Histone | -3.29 |
| AQK<sup>19</sup>NIAK | K<sup>35</sup>ESYAIYIK | N/A | A0A1S3DIC0_Histone | -3.15 |
| AVTK<sup>121</sup>YTSSK | ESYAIYIK<sup>44</sup>VLK | N/A | A0A1S3DIC0_Histone | -2.87 |
| K<sup>35</sup>ESYAIYIK | VLK<sup>47</sup>QVHPDTGVSSK | N/A | A0A1S3DIC0_Histone | -2.75 |
| AVTK<sup>121</sup>YTSSK | LLPGELAK<sup>109</sup>HAVSEGTK | N/A | A0A1S3DIC0_Histone | -2.21 |
| AVTK<sup>121</sup>YTSSK | VLK<sup>47</sup>QVHPDTGVSSK | N/A | A0A1S3DIC0_Histone | -2.1 |

<sup>1</sup> Crosslinked residue is indicated in bold with residue number in superscript following crosslinked lysine.
<sup>2</sup> Peptides containing PTMs are indicated with PTM type in subscript following modified lysine (Ac=acetylation; Me2=dimethylation).

**Figure 3.**
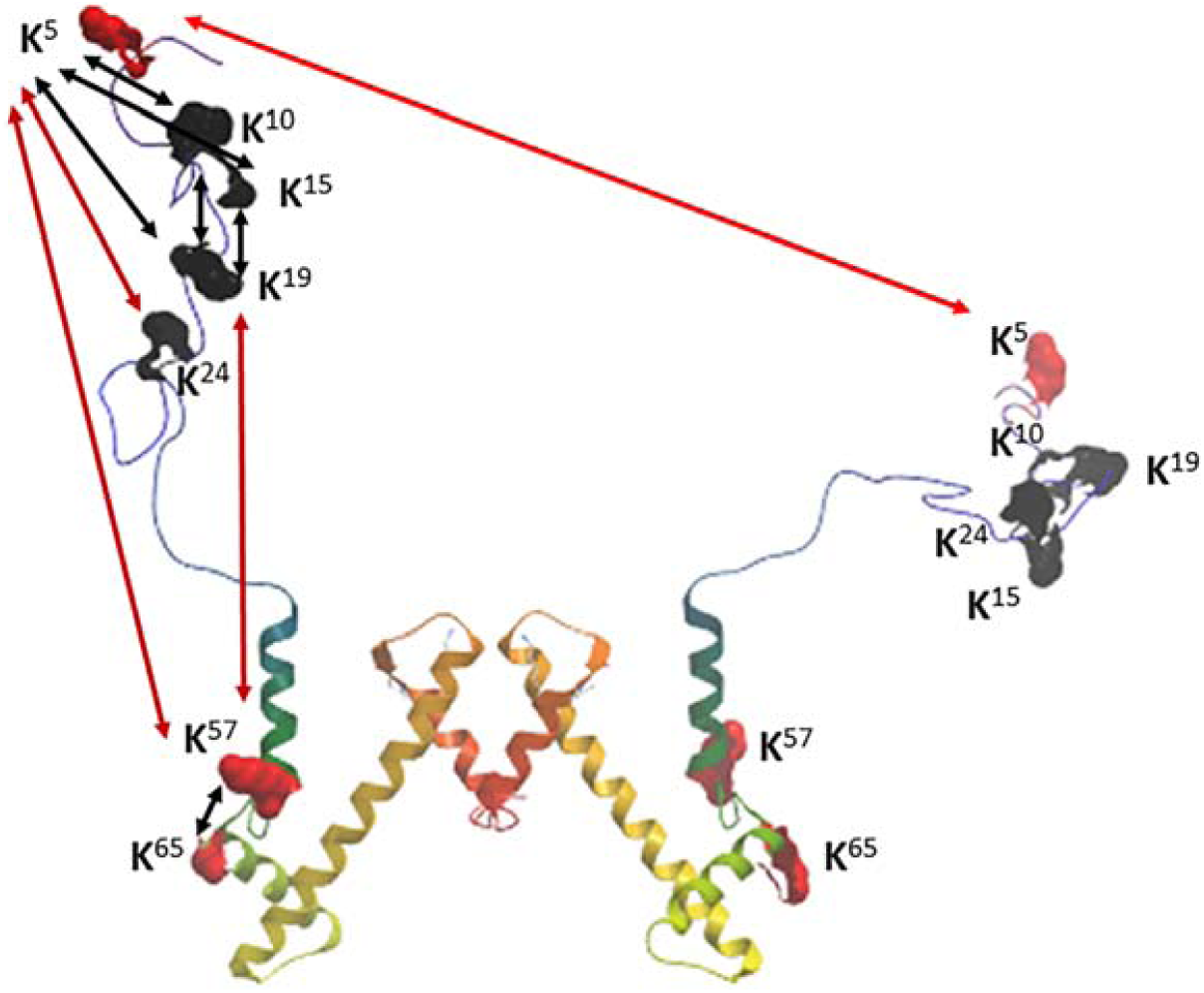
Structural representation of the two copies of histone H3 within a nucleosome highlighting residues involved in H3-H3 crosslinks found to be more abundant in nymph compared to adult insects. Distance between crosslinked residues less than 30 Å are represented by black lines, and distances greater than 30 Å are represented by red lines. The crosslinker size is 30 Å, and so the crosslinks represented by red lines are predicted to be intermolecular crosslinks between nucleosomes. Crosslinked lysine residues highlighted in black were found to have posttranslational modifications: acetylated at K^10^, K^15^, K^19^, K^24^, and dimethylated at K ^10^.

Four of the histone H3 crosslinks found to be more abundant in nymph compared to adult insects are predicted to be internucleosomal crosslinks, because the distance between crosslinked residues either within one H3 protein, or between the two H3 proteins within a nucleosome, is greater than the 30 Å crosslinker distance constraint (**Figure 3**).

Increased internucleosomal interactions in nymphs may be due to differences in chromatin configuration between nymph and adult insects. Alternative chromatin states have previously been identified between insect castes, wherein genetically identical organisms exhibit polyphenisms associated with distinct social identities within the colony (42, 43). Sequencing of chromatin immunoprecipitation (ChIP-seq) samples from worker and queen honeybee castes, and from major and minor worker carpenter castes, reveals caste-specific chromatin signatures associated with distinctive transcription profiles (44, 45). Nutritional differences between worker and queen bees have been associated with variation in histone PTMs, with a histone deacetylase inhibitor identified as a prominent component of royal jelly, the honeybee secretion which is consumed in greater abundance by queens compared to workers (46). Chromatin remodeling represents a effective method for an organism to effect the extensive and coordinated changes in gene and protein expression which characterize different insect developmental states. In previous studies of *C*Las(+) *D. citri*, significantly more nuclear fragmentation was observed in adult compared to nymph midgut cells, and protein biomarkers of apoptosis were more abundant in midguts of *C*Las(+) compared to *C*Las(-) adult insects (7, 47). Differences in histone crosslinks observed between *C*Las(+) nymph and adult *D. citri* in the present study may be related to the fact that while nuclear fragmentation and apoptosis were observed in adult midguts during *C*Las acquisition, nymph midgut nuclear morphology was largely unaffected by insect exposure to *C*Las.

Twenty-eight crosslinks involving metabolic proteins were upregulated in nymph compared to adult *D. citri* (**Table S3**). These include four intramolecular or dimeric crosslinks involving two aminopeptidase N proteins, two crosslinks between peptides from cathepsin proteases, and two crosslinks between peptides from carboxypeptidases. In related insect vectors, orthologs of these proteases have been found to play critical roles in pathogen transmission: a pea aphid (*Acyrthosiphon pisum*) aminopeptidase N was found to serve as the receptor mediating virus uptake into the gut (48), and a green peach aphid (*Myzus persicae*) cathepsin B was associated with reduced virus transmission (49). These highly abundant proteins are expressed in the gut epithelial cells of hemipterans and serve as sites of microbe-vector interaction at the molecular level.

A crosslink between hemocyanin 1 and hemocyanin 2 was identified as more abundant in nymph compared to adult *D. citri*. The hemocyanin 1 protein has previously been identified as a protein involved in the response of *D. citri* to *C*Las, and hemocyanin 2 has been found to be significantly more abundant in nymph compared to adult *D. citri (18)*. Hemocyanins are closely related to the larval storage proteins known as hexamerins, which have been characterized as playing crucial roles in insect development (22, 50). A related crosslink found to be more abundant in nymph *D. citri* was identified between two hexamerin proteins. Proteins with predicted roles in energy metabolism (*e*.*g*. ATP metabolism, electron transport in the mitochondria) were found to be part of crosslinks upregulated in both adult and nymph samples. In contrast, crosslinks between proteins involved in metabolism were identified as more abundant only in nymph samples.

A crosslink between two proteins annotated as Prohibitin-1 and Prohibitin-2 was found to be more abundant in *D. citri* nymph samples (**Table S3**). Prohibitins (PHBs) are pleitropic proteins localized to the inner mitochondrial membrane (51), and insect PHBs have been found to function in larval metabolism and pupal development in *Drosophila melanogaster* (52) and as a dengue virus receptor in *Aedes aegypti* (53). While no high-resolution crystal structural models for PHBs exist, a coiled-coil domain comprises the C terminal region of members of this protein family, which has previously been identified by PIR as a site of PHB homo- and hetero-dimerization in human cells (20, 54). There is substantial conservation between human and *D. citri* sequences, and strickingly, a *D. citri* PHB-1 homodimer was identified at the same residue (K202) as was found using PIR in human cells (20, 54). The PHB heterodimer which was found at higher levels in nymph compared to adults was represented by a crosslink between K202 of PHB-1 (the homodimer site), and K237 in the coiled-coil domain of PHB-2. A PHB heterodimer between these regions had also previously been identified in human cells. These data suggest a conserved function of the PHB interactions across the animal kingdom (20, 54).

### Posttranslational modifications of crosslinked peptides

Out of the 1571 unique peptide crosslinks identified in this study, 59 were found to contain at least one of the following PTMs: lysine acetylation (32), methylation (13), and dimethylation (14); and arginine acetylation (1) (**Table S4**). PTM residues are numbered in superscript and annotated in subscript (Ac=acetylation, Me=methylation, Me2=dimethylation, Me3=trimethylation). In four of these crosslinks, all mapping to the N terminal tail region of histone H3, PTMs were identified on both peptides in one crosslink. In three cases both peptides are acetylated, and in the fourth case one peptide is dimethylated (K^10^_Me2_STGG**K**^**15**^APR), and the other is acetylated (**K**^19^QLATK^24^_Ac_AAR) (**Table S4**). The dimethylated lysine on histone H3 aligns to the residue known as H3K9, and the acetylated lysine aligns to the residue known as H3K23. While H3K9 methylation (mono, di, and tri) has been associated with heterochromatin structures and decreased transcription (55, 56), H3K23 acetylation has been associated with transcriptional activation (57, 58). In 25 out of the 30 crosslinks containing acetylated lysines, both peptides mapped to one of four core histone proteins (histones H3, H4, H2A/2B). The five other acetylated crosslinked peptides mapped to a myosin heavy chain and an uncharacterized protein (**Table S4**). Three crosslinks were identified where both peptides map to the N terminal tail region of histone H3, with the same peptide found to be either acetylated and dimethylated at H3K9 (K^10^_Ac/Me2_STGG**K**^15^APR) in all three cases. An additional 24 crosslinks between peptides mapped to myosin or actin proteins were found to contain peptides with methylation or dimethylation PTMs.

### Protein homodimer crosslinks

Out of the 1245 crosslinks where each peptide in a pair is from the same protein, 65 crosslinks are considered to represent unambiguous homodimers due to the fact that the crosslinker-modified amino acid residue of each peptide is precisely the same (**Table S5**). Eight unique homodimer crosslinks were identified between peptides derived from the core histone proteins H4, H3.3, H2a, and H2B (**Table 4**).

**Table 4.** Crosslinked peptides representing unambiguous homodimers of histone proteins.

| Protein ID | Protein Description | Distance (Å)<sup>1</sup> | Peptide 1<sup>2</sup> | Peptide 2<sup>2</sup> |
| --- | --- | --- | --- | --- |
| MCOT08626.0.MT | Histone H2A | 14.2 | **K**<sup>62</sup>GNYAER | **K**<sup>62</sup>GNYAER |
| A0A1S3DIC0 | Histone H2B | 56.6 | AV**T****K**<sup>121</sup>YTSSK | AV**T****K**<sup>121</sup>YTSSK |
| MCOT05881.0.CC | Histone H3.3 | 116 | **T****K**<sup>5</sup>QTAR | **T****K**<sup>5</sup>QTAR |
| MCOT05881.0.CC | Histone H3.3 | 99.7 | **K**<sup>19</sup>QLAT**K**<sup>24</sup><sub>Ac</sub>AAR | **K**<sup>19</sup>QLAT**K**<sup>24</sup><sub>Ac</sub>AAR |
| MCOT05881.0.CC | Histone H3.3 | 15.6 | VTIMP**K**<sup>123</sup>DIQLAR | VTIMP**K**<sup>123</sup>DIQLAR |
| MCOT05881.0.CC | Histone H3.3 | 49.4 | Y**Q****K**<sup>57</sup>STELLIR | Y**Q****K**<sup>57</sup>STELLIR |
| MCOT08963.0.MT | Histone H4 | 13.5 | TVTAMDVVYAL**K**<sup>118</sup>R | TVTAMDVVYAL**K**<sup>118</sup>R |
| MCOT08963.0.MT | Histone H4 | 13.5 | TVTAMDVVYAL**K**<sup>118</sup>R | KTVTAMDVVYAL**K**<sup>118</sup>R |

<sup>1</sup> Distances in angstroms between homodimer residues of each protein pair within the nucleosome core particle predicted from the modeled X-ray structure. Distances greater than the 30 Å crosslinker length are predicted to represent interactions between nucleosomes, while shorter distances may represent interactions between protein pairs within a single histone octamer.
<sup>2</sup> Crosslinked lysine in bold and numbered in superscript, acetylated lysines indicated with K<sub>Ac</sub> and numbered in superscript.

The distance between two crosslinked **K**^123^ residues within a nucleosome is predicted to be 16 Å (**Table 4, Figure 4A)** and therefore could represent an intranucleosomal homodimer. This crosslinked residue is in the histone H3 C terminal region, which has been reported to be the H3 dimerization interface (59). In contrast, the predicted distance between two H3 N terminal residues (K19) within a nucleosome is 100 Å, suggesting that this crosslink represents an internucleosomal interaction (**Figure 4A**). Internucleosomal interactions have significant impacts on chromatin structure and gene transcription, are frequently mediated by posttranslational modifications of histone N terminal tail regions (60), and are indicative of chromatin in its closed (heterochromatin) rather than open (euchromatin) configuration (**Figure 4B**).

**Figure 4.**
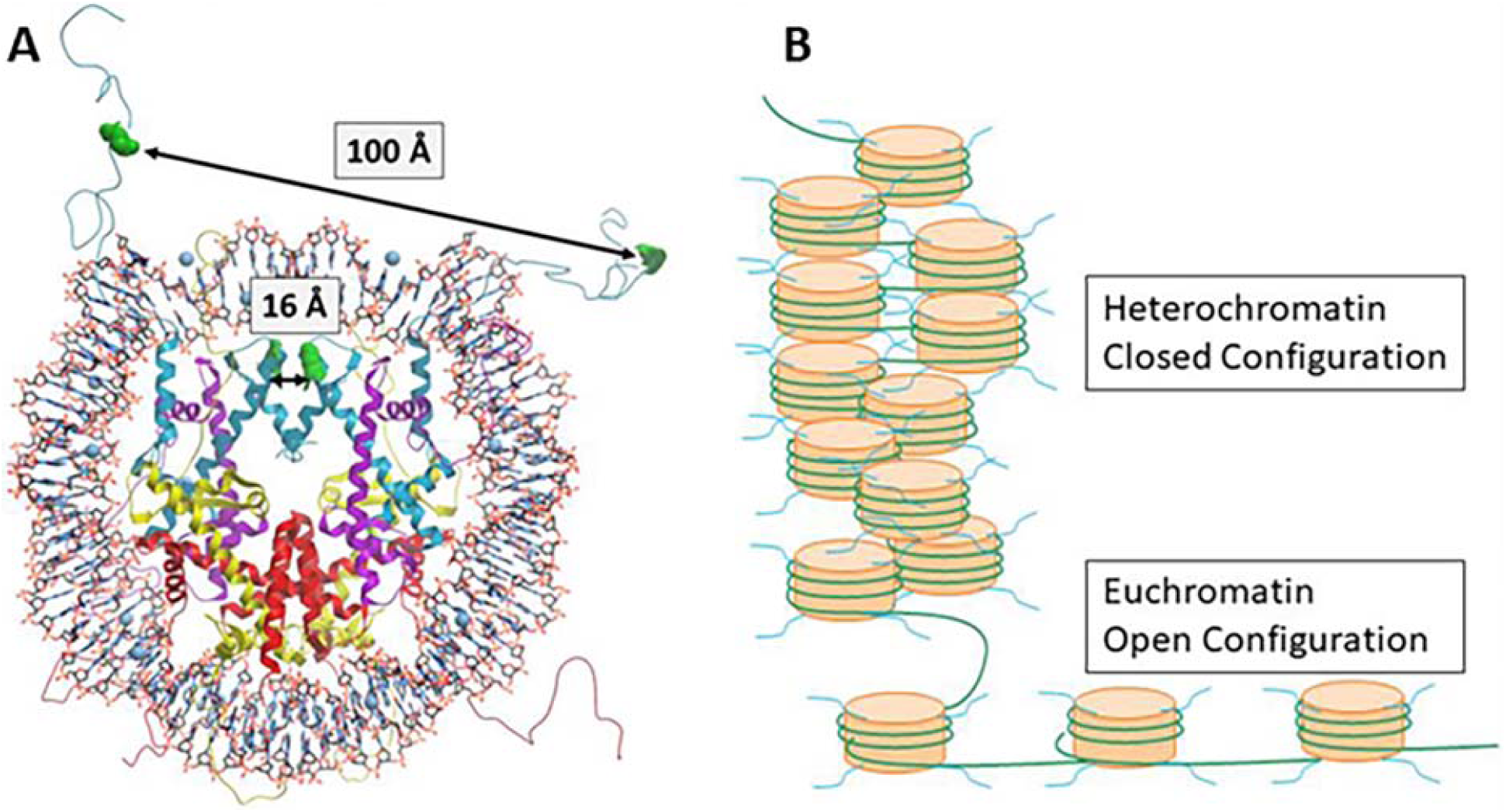
**a)** Structure of the nucleosome core particle as predicted from the X-ray diffraction pattern (PDB 1KX5). DNA is wound around the histone octamer, with histone N terminal tails extending from the nucleosome. The distances between the homodimer crosslink residues K123/K123 (16 Å) and K19/K19 (100 Å) in the two copies of histone H3 in the nucleosome is given. The distance between the two K19 residues is greater than the 30 Å crosslinker distance constraint, suggesting that the K19 homodimer represents an internucleosomal interaction between the H3 N terminal tails. **b)** Schematic of chromatin in its closed configuration (heterochromatin) and open configuration (euchromatin). DNA (green strand) is wrapped around nucleosome particles, with histone N terminal tails depicted in blue.

The H3K18 homodimer crossink residue is a well-characterized site for functionally significant histone acetylation – when acetylation at this site and others in histone H3 of a pathogenic fungus is blocked by a wheat microbiome bacteria, fungal growth and virulence is inhibited. Both peptides comprising this homodimer were acetylated at the lysine known as H3K23 (57, 58) (**K**^19^QLATK^24^_Ac_AAR), which is a histone acetylation associated with highly expressed genes in plants and insects (57, 58, 61) (**Table 4**). The tetramer comprising two H3/H4 dimers is predicted to form via interactions between C terminal regions of Histone H3 (62), which is represented in our data by the H3K123 homodimer (**Table 3, Figure 4A**). In addition, two unique peptide pairs were found to be crosslinked at the same lysine residue on the C terminus of Histone H4 (**Table 4**). The crosslinked residue corresponds to the Histone H4 C terminal lysine known as H4K91, which is found at the interface of the H3/H4 tetramer and H2A/H2B dimers and has been found to be acetylated and ubiqutinated with consequences on chromatin structure (63-65).

Other homodimers of biological significance which were identified in *D. citri* samples include three homodimer crosslinks between peptides in the C terminal domain of hemocyanin 1. Hemocyanin is known to form oligomeric structures, and has a predicted immunity function in numerous arthropod species (22, 50). The hemocyanin peptide LNH**K**^794^SFNYR, which was found in this study to form a homodimer, was previously identified as a component of a crosslink between hemocyanin and a *C*Las bacterial protein involved in coenzyme A biosynthesis (18).

### Bacterial endosymbiont protein crosslinks

Surprisingly, although the Percoll fractions were apparently enriched for CLas using qPCR detection, no crosslinks involving CLas proteins were identified. Eleven crosslinks involving peptides of bacterial endosymbionts of *D. citri* were identified – in one crosslink one peptide was derived from a bacterial protein and the other peptide was derived from an insect protein, while in the remainder both crosslinked peptides are bacterial (**Table 5**). The insect-bacterial crosslink involves a polyketide synthase (PedF) involved in production of diaphorin, an abundant polyketide produced by the *D. citri* endosymbiont ‘*Candidatus* Profftella armatura’, and an insect protein with a predicted secretion signal but no homology to known protein domains. Five of the bacterial crosslinks include Profftella proteins, including two crosslinks with both peptides mapping to a second polyketide synthase, PedI. The other five bacterial crosslinks involve the *D. citri* endosymbiont ‘*Candidatus* Carsonella rudii’, including a crosslink between the alpha and beta subunits of an ATP synthase, which are known interacting partners (66).

**Table 5.**
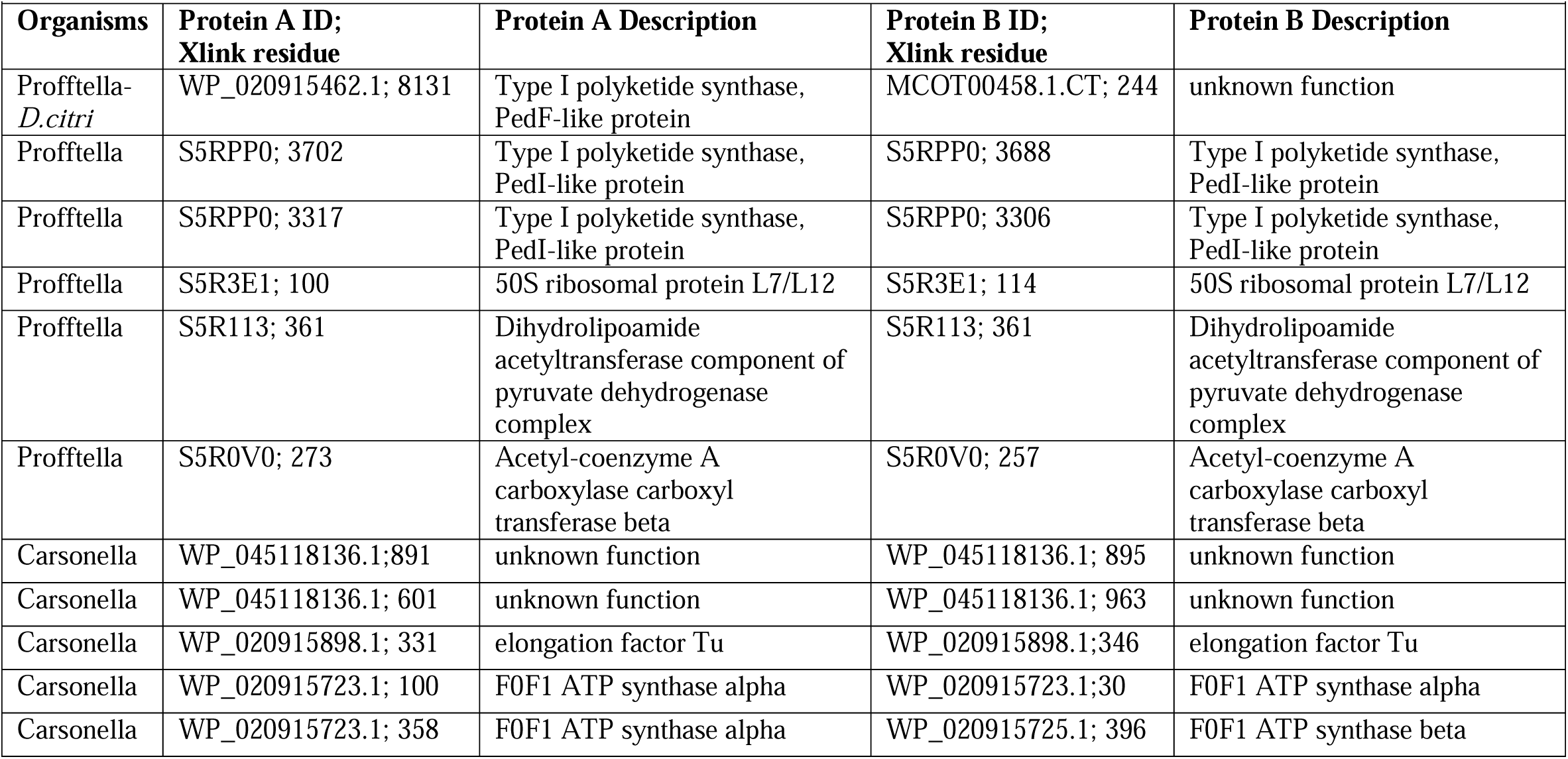
Crosslinks involving microbial proteins identified in *Diaphorina citri* samples.

| Organisms | Protein A ID;<br>Xlink residue | Protein A Description | Protein B ID;<br>Xlink residue | Protein B Description |
| --- | --- | --- | --- | --- |
| Profftella-<br>*D.citri* | WP_020915462.1; 8131 | Type I polyketide synthase, PedF-like protein | MCOT00458.1.CT; 244 | unknown function |
| Profftella | S5RPP0; 3702 | Type I polyketide synthase, PedI-like protein | S5RPP0; 3688 | Type I polyketide synthase, PedI-like protein |
| Profftella | S5RPP0; 3317 | Type I polyketide synthase, PedI-like protein | S5RPP0; 3306 | Type I polyketide synthase, PedI-like protein |
| Profftella | S5R3E1; 100 | 50S ribosomal protein L7/L12 | S5R3E1; 114 | 50S ribosomal protein L7/L12 |
| Profftella | S5R113; 361 | Dihydrolipoamide acetyltransferase component of pyruvate dehydrogenase complex | S5R113; 361 | Dihydrolipoamide acetyltransferase component of pyruvate dehydrogenase complex |
| Profftella | S5R0V0; 273 | Acetyl-coenzyme A carboxylase carboxyl transferase beta | S5R0V0; 257 | Acetyl-coenzyme A carboxylase carboxyl transferase beta |
| Carsonella | WP_045118136.1;891 | unknown function | WP_045118136.1; 895 | unknown function |
| Carsonella | WP_045118136.1; 601 | unknown function | WP_045118136.1; 963 | unknown function |
| Carsonella | WP_020915898.1; 331 | elongation factor Tu | WP_020915898.1;346 | elongation factor Tu |
| Carsonella | WP_020915723.1; 100 | F0F1 ATP synthase alpha | WP_020915723.1;30 | F0F1 ATP synthase alpha |
| Carsonella | WP_020915723.1; 358 | F0F1 ATP synthase alpha | WP_020915725.1; 396 | F0F1 ATP synthase beta |

### Quantitative mass spectrometry proteomics of D. citri percoll fraction samples

Adult and nymph *Diaphorina citri* insects were collected from healthy (*C*Las-) and infected (*C*Las+) citrus plants. Percoll gradient fractionation of insect homogenate was used to generate a concentrated sample of insect and microbe cells. The same fractions from the percoll gradient as were used for PIR analysis - those selected as having the highest abundance of *C*Las based on qPCR analysis for *C*Las titer - were used as starting material for quantitative proteomcis. Proteins were extracted from percoll fraction samples, and following sample preparation, trypsin-digested peptides were analyzed by mass spectrometry. A total of 1780 insect and microbe proteins were identified (**Table S6**). More than 98% of these proteins are insect proteins, and 37 microbe proteins derived from the three primary bacterial endosymbionts of *D. citri* [‘*Candidatus* Profftella armatura’ (22), *Wolbachia pipientis* (11), and ‘*Candidatus* Carsonella rudii’ (4)] were identified. Peptides were also assigned to two *C*Las ATPase proteins (WP_015452542.1, WP_015452544.1) and a *C*Las DnaK molecular chaperone (WP_015824904.1). However, these peptide spectra were identified in both *C*Las(-) and *C*Las(+) samples, and the protein region to which the peptides map is highly conserved between *C*Las and the three endosymbionts, suggesting that these proteins are likely derived from one of the endosymbionts, which are present in all sample types.

### Statistical analysis of quantitative mass spectrometry data

Statistical analysis of spectral counts was performed to identify proteins which were differentially abundant between sample categories (healthy nymph vs healthy adult, infected nymph vs infected adult, healthy nymph vs infected nymph, healthy adult vs infected adult, **Table 6, Table S7**). The fewest number of differentially abundant proteins (50) were identified between *C*Las(-)/*C*Las(+) nymphs, whereas nearly three times as many differentially abundant proteins (132) were identified between *C*Las(-)/*C*Las(+) adults (**Table 6**). These results are consistent with data from a previous quantitative proteomics analysis of whole insect nymph and adult samples – in this prior study, four times as many proteins in the adult insect were regulated by *C*Las infection as in the nymph insect (18). Many more proteins (>600) were found at significantly different abundance between nymph and adult percoll fraction samples than between *C*Las(+) and *C*Las(-) samples (**Table 6, Table S7**). Vitellogenins are proteins with documented function in lipid transport, reproduction, and immunity in insects. While some vitellogenins which function in reproduction are expressed predominantly in female insects, a vitellogenin in the potato psyllid (*Bactericera cockerelii)* was found to be expressed at similar levels in male and female insects and may have an immune function (67). Several vitellogenin proteins, which were found at higher levels in this study in *C*Las(+) adult compared to *C*Las(-) adult samples, were either undetected or found at the limit of detection in nymphs of both infection status (**Table S7**). These include Vg-1 vitellogenins (XP_008476565, MCOT18962.0.CT), which were previously identified in the hemolymph of adult *D. citri* both as among the most highly abundant proteins in this tissue, and as more abundant in hemolymph from *C*Las(+) compared to *C*Las(-) insects (13).

**Table 6.** Summary of statistical analysis of quantitative proteomics data.

| Sample Category Comparison (A vs B) | Number upregulated proteins<sup>1</sup> | Number downregulated proteins<sup>1</sup> |
| --- | --- | --- |
| CLas(-) adult vs CLas(-) nymph | 336 | 352 |
| CLas(+) adult vs CLas(+) nymph | 294 | 303 |
| CLas(+) adult vs CLas(-) adult | 65 | 67 |
| CLas(+) nymph vs CLas(-) nymph | 25 | 25 |

<sup>1</sup> For each sample category comparison, the number of proteins upregulated and downregulated in category A vs B are given.

Subsequent to performing this work, data from our lab has revealed substantial variability between the different color morphs which are found within a population of *D. citri* (68). Because blue and non-blue *D. citri* were pooled into the same sample in this study, the abundance of proteins which may be significantly different between color morphs will not be precisely quantified. Follow-up work to separate insects from each developmental stage into separate color morph classes for analysis will provide insight into the impact of this significant polyphenism on insect development and immunity.

### Integration of PIR and Spectral Count data

For the peptide crosslinks identified at significantly higher levels in nymph or adult *D. citri*, the increased abundance of these crosslinks may be attributed to the higher abundance of the parent proteins in one developmental stage relative to the other. However, in several instances a crosslink was found at significantly higher levels in nymphs compared to adults, but the proteins from which the peptides comprising that crosslink were derived were found at significantly lower levels in nymphs compared to adults. The PHB1-PHB2 heterodimer crosslink was found at four-fold higher abundance in nymphs compared to adults, but the spectral count data indicate that PHB1 and PHB2 are found at nine-fold and five-fold higher levels, respectively, in adults compared to nymphs. This indicates that protein complex formation is not simply a function of the concentration of the constituent proteins, and suggests that differences in protein complex and/or protein interaction topology is responsible for differences in crosslink abundance. Additional proteins involved in promoting PHB1-PHB2 complex formation may be more abundant in nymphs relative to nymphs, leading to increased abundance of this crosslink in nymphs despite the decreased abundance of the component proteins. Similarly, a crosslink between two hemocyanin 1 peptides was found by PIR to be more abundant in *C*Las(+) nymphs compared to adults, while spectral counting data indicate that hemocyanin 1 is one of the most abundant proteins in *C*Las(+) adults, and is over ten times as abundant in these samples compared to *C*Las(+) nymphs (**Tables S3, S7**).

Statistical analysis of spectral counting data suggests that histone protein expression is regulated by insect developmental state and *C*Las infection status. Histone H2B (two isoforms) and histone H4 were both found at higher abundance in adult compared to nymph insects, regardless of infection status (**Table 7**). In contrast, multiple crosslinks involving these core histones were found to be more abundant in nymphs compared to adults (**Tables 2, 3**). *C*Las infection is associated with reduced histone H2B expression: both histone H2B isoforms were found at higher abundance in *C*Las(-) compared to *C*Las(+) nymph and adult insects. Histone H4 was also more abundant in *C*Las(-) compared to *C*Las(+) adult insects, but no statistical difference between spectral counts was found between *C*Las(-) and *C*Las(+) nymph samples. The core histone variant H3.3 was found at higher levels in *C*Las(-) nymphs compared to *C*Las(-) adults, but in *C*Las(+) samples the protein was more abundant in adults compared to nymphs (**Table 7**).

**Table 7.** Histone proteins found to be differentially abundant between insect sample classes based on development stage and *C*Las infection status.

| Protein ID | Annotation | N<sup>(-)</sup> | N<sup>(+)</sup> | A<sup>(-)</sup> | A<sup>(+)</sup> | Significant Difference<sup>1</sup> |
| --- | --- | --- | --- | --- | --- | --- |
| MCOT10597.0.OO | Histone H2B | 80.49 | 15.35 | 129.08 | 43.66 | Development: N<sup>(-)</sup><A<sup>(-)</sup>; N<sup>(+)</sup><A<sup>(+)</sup>;<br>CLas infection: N<sup>(+)</sup><N<sup>(-)</sup>; A<sup>(+)</sup><A<sup>(-)</sup> |
| A0A1S3DIC0 | Histone H2B | 199.48 | 61.42 | 294.43 | 134.16 | Development: N<sup>(-)</sup><A<sup>(-)</sup>; N<sup>(+)</sup><A<sup>(+)</sup>;<br>CLas infection: N<sup>(+)</sup><N<sup>(-)</sup>; A<sup>(+)</sup><A<sup>(-)</sup> |
| MCOT08963.0.MT | Histone H4 | 63.33 | 44 | 78 | 63.33 | Development: N<sup>(-)</sup><A<sup>(-)</sup>; N<sup>(+)</sup><A<sup>(+)</sup>;<br>CLas infection: A<sup>(+)</sup><A<sup>(-)</sup> |
| MCOT05881.0.CC | Histone H3.3 | 17.99 | 6.22 | 13.16 | 11.56 | Development: N<sup>(-)</sup>>A<sup>(-)</sup>; N<sup>(+)</sup><A<sup>(+)</sup> |
| XP_008482743.1 | Histone H1E | 9.01 | 3.95 | 3.14 | 2.33 | Development: N<sup>(-)</sup>>A<sup>(-)</sup>; N<sup>(+)</sup>>A<sup>(+)</sup> |
| XP_008480034.1 | Histone H1.M6.2 | 36 | 12.66 | 4.66 | 3.33 | Development: N<sup>(-)</sup>>A<sup>(-)</sup>; N<sup>(+)</sup>>A<sup>(+)</sup> |
| MCOT07573.0.MM | Histone H1-II-1 | 161.86 | 67.86 | 50.76 | 40.93 | Development: N<sup>(-)</sup>>A<sup>(-)</sup>; N<sup>(+)</sup>>A<sup>(+)</sup>;<br>CLas infection: N<sup>(+)</sup><N<sup>(-)</sup>; A<sup>(+)</sup><A<sup>(-)</sup> |
| MCOT12721.3.CO | Micronuclear<br>Histone H1 | 11.9 | 1.65 | 0.66 | 0.32 | Development: N<sup>(-)</sup>>A<sup>(-)</sup>; N<sup>(+)</sup>>A<sup>(+)</sup>;<br>CLas infection: N<sup>(+)</sup><N<sup>(-)</sup> |

<sup>1</sup> Average spectral counts for each sample class (N=3), and sample classes found to have significantly different spectral counts, are listed. N<sup>(-)</sup>=nymph CLas(-); N<sup>(+)</sup>=nymph CLas(+); A<sup>(-)</sup>=adult CLas(-); A<sup>(+)</sup>=adult CLas(+).

Whereas histone H2B, H4, and H3.3 are all components of the nucleosomal core particle around which DNA are wound, linker histone H1 proteins bind to the DNA entry and exit sites on the nucleosome, stabilizing chromatin structure and influencing gene expression (69, 70). In contrast to the higher expression of histone H2B and H4 in adults compared to nymphs, four histone H1 variants were found to be much more abundant in nymph compared to adult insects regardless of infection status. In addition, micronuclear histone H1 (nymphs) and histone H1-II-1 (adults and nymphs) were also more abundant in *C*Las(-) compared to *C*Las(+) insects (**Table 7**). Two proteins which function in histone binding and modification were found at significantly different levels between sample classes: a histone lysine N-methyltransferase (MCOT17739.0.CT) was significantly more abundant in *C*Las(+) compared to *C*Las(-) adult samples, while the histone binding protein rbbp4 (MCOT00465.0.CT) was significantly more abundant in *C*Las(+) adult compared to *C*Las(+) nymph samples (**Table S7**).

### Identification of posttranslational modifications in quantitative proteomics data

Lysine acetylation, methylation, and dimethylation were selected as variable modifications during Mascot searching. The data are summarized as the number of assigned modified spectra per protein (**Table S8**), and the modified peptide sequences are available in the peptide spectrum report (**Table S9**). Peptides with acetylated lysines were mapped to 112 proteins, peptides with methylated lysines were mapped to 255 proteins, and peptides with dimethylated lysines were mapped to 103 proteins (**Table S8**). Acetylated and methylated peptides were mapped to several of the histone proteins which were found at different abundance in adult compared to nymph and/or *C*Las(-) compared to *C*Las(+) samples (**Tables 7**,**8**). The acetylated peptide K^82^_Ac_YLVSAVAAGK is found in two histone H1 proteins found to be more abundant in nymph compared to adult samples. Based on our data it is not possible to definitively map this acetylated peptide to either of these proteins, or to know whether both of these proteins, or only one of them, are acetylated at this peptide. The acetylated residue is found at K82 on XP_008482743.1 (histone H1E) and K88 on MCOT07573.0.MM (histone H1-II-1). This acetylated peptide aligns to K85 of the human histone H1 (AAA63187.1), which is subject to post-translational acetylation significantly impacting chromatin structure (71). Both of these histone H1 proteins were more abundant in nymph compared to adult samples (**Table 7**). While the peptide KYLVSAVAAGK is found in both adult and nymph samples, the modified peptide with the first lysine acetylated is only found in nymph samples (**Table 8**).

**Table 8.** Acetylated and methylated peptides mapping to the histone proteins identified by quantitative proteomics as differentially abundant between *D. citri* sample classes.

|  |  |  | # modified/unmodified peptide spectra<sup>1</sup> |  |  |  |
| --- | --- | --- | --- | --- | --- | --- |
| Protein ID | Protein Description | Modified Peptide<sup>2</sup> | nymph CLas(-) | nymph CLas(+) | adult CLas(-) | adult CLas(+) |
| XP_008482743.1 | histone H1E | K<sup>82</sup><sub>Ac</sub>YLVSAVAAGK* | 12/93 | 6/39 | 0/36 | 0/39 |
| XP_008480034.1 | histone H1.M6.2 | KPAQAQPAAGK<sup>59</sup><sub>Me</sub> | 24/0 | 1/0 | 6/0 | 5/0 |
| A0A1S3DIC0 | histone H2B | QVHPDTGVSSK<sup>58</sup><sub>Ac</sub>* | 14/88 | 6/36 | 0/66 | 4/68 |
| A0A1S3DIC0 | histone H2B | QVHPDTGVSSK<sup>58</sup><sub>Me</sub>* | 4/88 | 6/36 | 0/66 | 0/68 |
| A0A1S3DIC0 | histone H2B | VLK<sup>47</sup><sub>Ac</sub>QVHPDTGVSSK* | 0/62 | 0/32 | 2/40 | 0/28 |
| A0A1S3DIC0 | histone H2B | HAVSEGTK<sup>117</sup><sub>Me</sub> | 10/657 | 5/119 | 10/1062 | 12/346 |
| A0A1S3DIC0 | histone H2B | LLPGELAK<sup>109</sup><sub>Me</sub> | 20/64 | 6/26 | 42/67 | 47/63 |
| MCOT05881.0.CC | Histone H3.3 | K<sup>28</sup><sub>Ac</sub>SAPSTGGVK | 1/9 | 3/2 | 0/4 | 0/3 |
| MCOT05881.0.CC | Histone H3.3 | K<sup>28</sup><sub>Me2</sub>SAPSTGGVK | 20/9 | 27/2 | 17/4 | 16/3 |
| MCOT05881.0.CC | Histone H3.3 | STGGK<sup>15</sup><sub>Ac</sub>APR | 44/0 | 12/0 | 22/0 | 10/0 |
| MCOT05881.0.CC | Histone H3.3 | QLATK<sup>24</sup><sub>Ac</sub>AAR* | 44/0 | 8/0 | 25/0 | 28/0 |
| MCOT05881.0.CC | Histone H3.3 | EIAQDFK<sup>80</sup><sub>Me2</sub>TDLR* | 4/42 | 4/22 | 22/36 | 14/40 |
| MCOT05881.0.CC | Histone H3.3 | EIAQDFK<sup>80</sup><sub>Me</sub>TDLR* | 34/42 | 14/22 | 68/36 | 66/40 |
| MCOT08963.0.MT | Histone H4 | DAVITYTEHAK<sup>104</sup><sub>Me</sub> | 2/31 | 1/8 | 0/30 | 0/33 |
| * These peptides each map to 2 unique proteins (XP_008482743.1/MCOT07573.0.MM; XP_008481587.1/MCOT05138.0.CC; MCOT05881.0.CC/XP_008484413.1) and cannot be assigned to one or the other based on these data |  |  |  |  |  |  |

<sup>1</sup> The total number of modified/unmodified spectra identified in each sample class (sum of three reps/class) are given.
<sup>2</sup> Modification type and residue number are indicated for each modified lysine.

Multiple methylated and acetylated peptides derived from histone H2B, H3.3, and H4 were identified (**Table 8**). Unique methylated and acetylated forms of a single histone H2B peptide were among five modified peptides mapping to histone H2B. Six modified peptides mapped to histone H3.3, including two acetylated peptides (STGGK^15^_Ac_APR, QLATK^24^_Ac_AAR) for which no spectra representing unmodified peptides were identified (**Table 8**). These acetylated peptides, for which more spectra were found in nymph *C*Las(-) samples compared to all other sample types, were identified as part of five separate crosslinks found by PIR to be more abundant in nymph *D. citri* (**Figure 5**). The acetylated peptides identified by PIR are identical except for the presence of an additional N terminal lysine, which is the crosslinked residue (**K**^10^STGGK^15^_Ac_APR, **K**^19^QLATK^24^_Ac_AAR).

**Figure 5.**
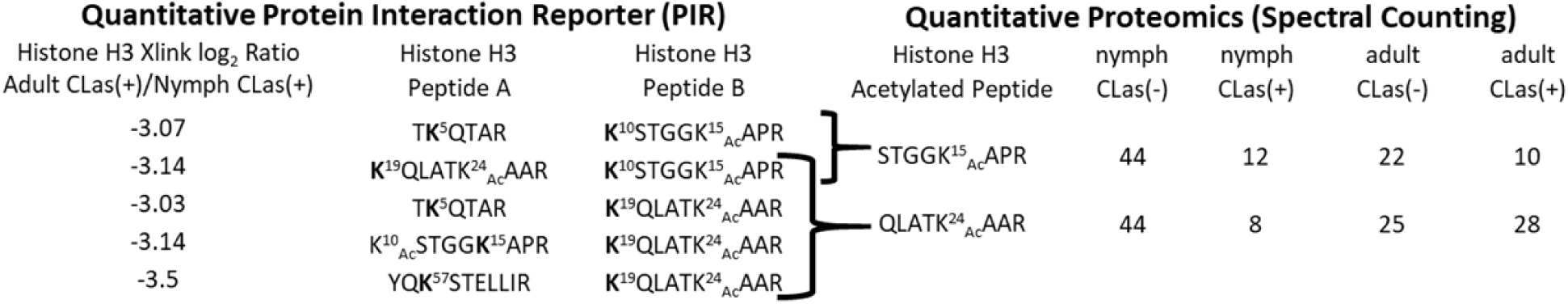
Impact of development and CLas infection on histone H3 acetylation: integration of PIR and spectral counting data. PIR crosslinks found to be more abundant in nymph compared to adult *D. citri* involve lysine-acetylated peptides (K15, 24) in the N terminal tail region histone H3. Crosslinked residues are in bold. Peptides containing these acetylated lysines were also identified in spectral counting data obtained from separate samples. Total spectral counts for the two modified peptides in the four sample classes (sum of three reps/class) are given.

The presence of an additional lysine on the peptides identified by PIR can be explained by the presence in the histone H3 protein sequence of an arginine residue on the N terminal side of this lysine. The trypsin enzyme used in peptide sample preparation for both PIR and spectral counting analysis cuts on the carboxyl side of either a lysine or arginine. In the PIR samples it is possible that the presence of the crosslinker on the lysine residue influences trypsin digestion such that the preferred cutting site is on the carboxyl side of the upstream arginine. In this way the N terminal lysine is retained in the PIR samples, while in the peptides prepared for quantitative proteomics the preferred trypsin digestion site is on the carboxyl side of this lysine.

## Conclusions

Complementary proteomics analyses were used to identify and quantify proteins and protein interactions in nymph and adult samples of *Diaphorina citri*, the insect vector of the bacterial pathogen (‘*Candidatus* Liberibacter asiaticus’, *C*Las) associated with citrus greening disease. Quantitative isotope-labeled protein interaction reporter (PIR) technology identified more than 1500 unique crosslinks between pairs of insect peptides, between pairs of peptides derived from insect bacterial endosymbionts, and in one case between an insect and a microbe peptide. Crosslinker data analysis revealed that insect histone protein crosslinks were overrepresented among three classes of protein interactions: unambiguous homodimers, crosslinks more abundant in nymph compared to adult insects, and crosslinks containing methylated and acetylated lysines. Quantitative proteomics spectral counting analysis of nymph and adult insects collected from healthy and infected citrus trees [*C*Las(-) and *C*Las(+) samples] indicated that development and infection status influence histone protein levels and posttranslational modification of histone peptides. While histone H2B and H4 were both found at higher abundance in adult compared to nymph insects, multiple crosslinks involving these core histones were found to be more abundant in nymphs compared to adults. These histone crosslinks may represent PTM-mediated internucleosomal interactions reflecting developmental stage-specific chromatin configurations. Several crosslinked peptides identified in psyllids have been found in crosslinked peptides derived from human cells, showing selection for particular protein interaction topologies between orthologous proteins conserved across the animal kingdom. Variation in histone protein abundance, protein interactions, and postranslational modifications between nymph and adult insects may reflect changes in chromatin structure during development with potentially dramatic consequences on transcriptional activity and vector competence of this globally significant agricultural pest.

## Data Availability

The mass spectrometry proteomics crosslinking data have been deposited to the ProteomeXchange Consortium via the PRIDE partner repository with the dataset identifier PXD023382. The mass spectrometry proteomics spectral counting data have been deposited to the ProteomeXchange Consortium via the PRIDE partner repository with the dataset identifier PXD020412.

## Supplemental Data

This article contains supplemental data.

